# Muscle group specific transcriptomic and DNA methylation differences related to developmental patterning in FSHD

**DOI:** 10.1101/2021.09.28.462147

**Authors:** Katherine Williams, Xiangduo Kong, Nam Viet Nguyen, Cassandra McGill, Rabi Tawil, Kyoko Yokomori, Ali Mortazavi

## Abstract

Muscle groups throughout the body are specialized in function and are specified during development by position specific gene regulatory networks. In developed tissue, myopathies affect muscle groups differently. Facioscapulohumeral muscular dystrophy, FSHD, affects upper body and tibialis anterior (TA) muscles earlier and more severely than others such as quadriceps. To investigate an epigenetic basis for susceptibility of certain muscle groups to disease, we perform DNA methylation and RNA sequencing on primary patient derived myoblasts from TA and quadricep for both control and FSHD2 as well as RNA-seq for myoblasts from FSHD1 deltoid, bicep and TA over a time course of differentiation. We find that TA and quadricep retain methylation and expression differences in transcription factors that are key to muscle group specification during embryogenesis. FSHD2 patients have differences in DNA methylation and expression related to SMCHD1 mutations and FGF signaling. Genes induced specifically in FSHD are more highly expressed in commonly affected muscle groups. We find a set of genes that distinguish more susceptible muscle groups including development-associated TFs and genes involved in WNT signaling. Adult muscle groups therefore retain transcriptional and DNA methylation differences associated with development, which may contribute to susceptibility in FSHD.

## Background

The human body has over 650 named skeletal muscle groups [1,2]. These groups are heterogeneous in terms of compositions of fast and slow twitch fibers and regenerative capabilities [3]. Initial muscle cell specification through the activation of the key myogenic factors MRF4/MYF5, MYOD, and MYOG is regulated by upstream transcription factors that vary depending on the location of the muscle cells [4,5]. PITX2 plays a central role in regulating MYF5/MRF4 in extraocular muscles and can regulate MYOD in limb muscles [4–7]. SIX and EYA TFs activate PAX3 and MYF5/MRF4 in limb muscles [7].

The limbs are initially specified by HOX genes which activate TBX5 in the forelimb and PITX1 in the hindlimb [8]. PITX1 then activates TBX4 in the hindlimb [8]. TBX5 and TBX4 activate FGF10 which forms a gradient with FGF8 that is expressed at the apical ectodermal ridge to control limb outgrowth [8]. Limb outgrowth specifies the proximal/distal axis resulting in three major regions; the stylopod, zeugopod and autopod [8]. MEIS factors are expressed proximally in the stylopod but are repressed by SHOX2 distally into the zeugopod where HOXA11 is expressed [9,10]. Dorsal/ventral patterning is controlled by WNT and LMX1B expression on the dorsal side and EN1 on the ventral side which represses WNT [11].

Gene expression in embryonic development of different muscle groups is well studied, but few studies have surveyed gene expression in adult tissue [12,13]. An estimated 50% of transcripts are differentially expressed in adult muscle with some heterogeneity coming from fiber-type composition and expression of developmental related genes [14]. In addition to transcription, adult muscle groups retain DNA methylation differences [15]. The molecular differences between muscle groups may contribute to severity of affectedness in myopathies.

Many myopathies affect muscle groups of the body exclusively or more severely, such as facioscapulohumeral muscular dystrophy (FSHD), which is characterized by noticeable weakness in the most commonly affected muscle groups in the upper body including facial and humeral [16,17]. FSHD progression into muscle groups is sporadic, but some groups such as the tibialis anterior are more commonly affected or affected earlier [18]. Certain muscle groups are less frequently affected or affected later including the quadricep and the deltoid [19,20].

FSHD is caused by the misexpression of the embryonic transcription factor *DUX4* in skeletal muscle [17,19]. DUX4 activates expression of its target genes including a number of embryonic related transcription factors and chromatin remodelers as well as repeat elements including endogenous retroviruses such as ERVLs [21–24]. DUX4 target gene expression correlates with signs of active disease [25,26]. Comparison of the commonly affected bicep to the less affected deltoid found greater expression differences between FSHD and control in bicep than deltoid but did not assess differences in DUX4 target gene expression between the two muscle groups [27].

We survey transcription and DNA methylation to establish differences between muscle groups that may contribute to susceptibility in FSHD. We perform RNA-seq and probe enriched bisulfite sequencing to survey transcriptional and DNA methylation differences between tibialis anterior (TA) and quadricep in myoblasts from healthy patients and those with FSHD2 with SMCHD1 mutations. We examine differences in DNA methylation and expression between TA and quadricep, and find retained differences in TFs important for development. Next, we find a set of genes specifically upregulated in FSHD2 over time including DUX4 target genes. Notably, the promoters of these genes are highly methylated in both FSHD2 and control cells. We look at genome-wide DNA methylation and gene expression differences in FSHD2 and find hypomethylation and increased expression for genes in regions regulated by SMCHD1. We survey differences for TE loci and find repeat elements upregulated in response to DUX4 are especially upregulated in FSHD2 cells from TA but not quadricep. To determine possible differences in gene expression correlating with susceptibility, we performed RNA-seq for myoblasts from the TA, bicep and deltoid of FSHD1 patients. We identify muscle group specific gene expression for TA, quadricep, bicep and deltoid, and genes differentially expressed between muscle groups with different susceptibilities to FSHD.

## Results

### Muscle group specific DNA methylation and gene expression in developmental TFs

To identify epigenetic differences between muscle groups, we performed capture enrichment bisulfite sequencing (Methods) on two primary myoblast cell lines from quadricep and two from tibialis anterior (TA) for days 0, 3 and 12 of differentiation (Figure S1). We merged CpG sites within 200 bp into regions which were then filtered for coverage (Methods). We found very few differentially methylated regions (DMRs) (percent diff 25, q-value <0.01) between the differentiation days and decided to treat the different days as replicates for further comparisons (Figure S2A, S2B).

We compared TA and quadricep and found 1081 regions more highly methylated in quadricep as well as 686 regions with higher methylation in TA (Figure 1A, S2B). The regions with the highest percent methylation differences are associated with transcription factors that play a role in limb specific muscle specification. These include *PITX2*, which controls location specific gene networks to induce MRFs [4]. Also included is *MEIS1*, which is expressed in the stylopod and is more highly methylated in TA than quad (Figure 1A) [9]. Of the 1767 DMRs, 357 are associated with 265 TFs, and 83 of those are involved in pattern specification while 39 are involved in muscle structure development (Figure S2C, Table S1). In support of this, gene ontology analysis for these DMRs revealed significant association with development and structure morphogenesis (Figure 1B). Quadricep and TA therefore retain DNA methylation patterns from their specification.

**Figure 1:**
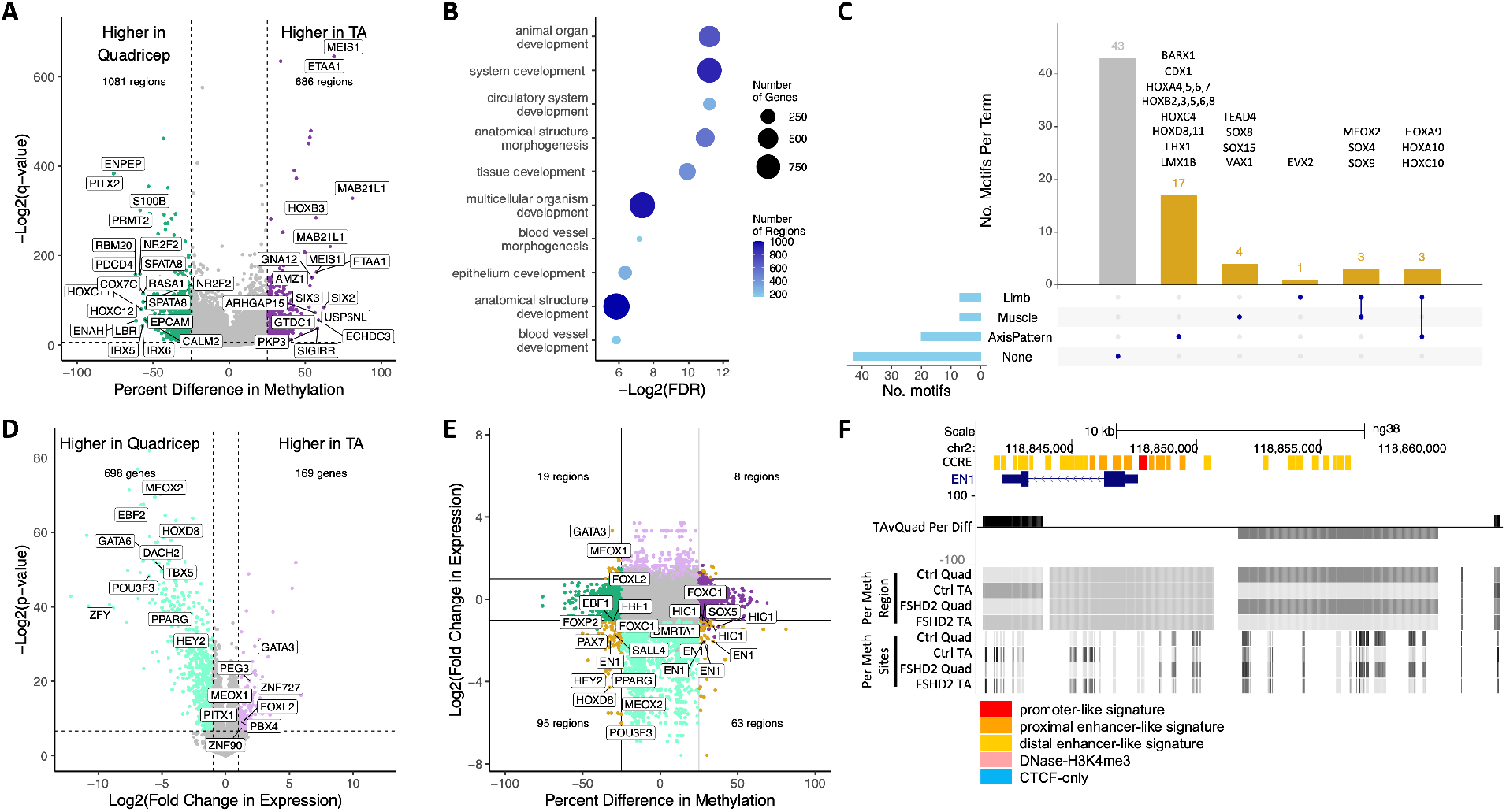
Adult muscle cells retain differences in DNA methylation and gene expression for transcription factors involved in developmental patterning. **(A)** Volcano plot of DNA methylation differences for control cells from TA versus quadricep. Regions with a p-value of >0.01 or a percent methylation difference of <25 are taken as not significant and colored in grey. Regions with more methylation in TA are in purple. Regions with more methylation in quadricep are in green. The top 10 regions with the highest percent difference in the positive or negative directions are labelled. The numbers of regions higher in TA and quadricep are labelled at the top. Vertical lines intersect y-axis at -25 and 25 percent. Horizontal line intersects y-axis at - log2(0.01). **(B)** Gene ontology terms enriched in differentially methylated regions between TA and quadricep. The top 10 significant terms (p-value <0.05) with at least 10 genes associated to the term from the background set of regions detected are shown. Color indicates number of differentially methylated regions in the term. Size indicates the number of genes associated with the differentially methylated regions in the term. **(C)** Upset plot of gene ontology terms of transcription factors whose motifs are enriched in differentially methylated regions between TA and quadricep. Each transcription factor was annotated with GO terms related to limb, muscle or axis patterning (see Methods for details). The number of transcription factors found enriched in the DMRs that falls into the individual categories are shown in blue on the left. The number of transcription factors belonging to the interest of the categories indicated in dark blue are in gold and grey on the top. **(D)** Volcano plot of gene expression differences for control cells from TA versus quadricep. Genes with a p-value of >0.01 or an absolute log2(fold change in expression) <1 are in grey. Genes with higher expression in the TA are in light purple. Genes with higher expression in the quadricep are in light green. The numbers of genes with higher expression in TA or quadricep are labelled at the top. The 10 transcription factors with the lowest p-values for positive or negative fold change are labelled. Vertical lines intersect y-axis at -1 and 1 log2(fold change). Horizontal line intersects y-axis at -log2(0.01). **(E)** Scatterplot of differences in DNA methylation and expression of nearby gene for TA versus quadricep. Regions are associated with genes through GREAT. Regions with significant differences in both DNA methylation and gene expression are in gold, and the numbers of these regions are labelled in the corresponding quadrant. All regions associated with transcription factors in the given quadrants are labelled. Regions associated with genes with significant differences in gene expression but not DNA methylation are colored in light purple and light green. Regions with significant differences in DNA methylation but not in expression of the associated gene are colored in green and purple. Vertical lines intersect x-axis at -25 and 25 percent. Horizontal lines intersect y-axis at -1 and 1 log2(fold change). **(F)** UCSC genome browser shot of DNA methylation near *EN1. EN1* gene model is from gencode v28. Percent methylation for each group of samples in the merged region are labelled with increase in methylation indicated with darker grey. The percent difference in methylation is indicated as a barplot with higher methylation in TA as positive and higher methylation in quadricep as negative. Percent methylation at individual sites for the given groups are at the bottom with darker color indicating higher methylation.

We looked for TF motifs enriched in the DMRs between TA and quadricep to see what could be regulating these regions. The DMRs are enriched for motifs of developmental transcription factors (Figure S2D). Importantly, we found motifs for key transcription factors involved in axis specification, skeletal muscle development and differentiation, and limb specification and morphogenesis, including motifs for 19 TFs controlling anterior/posterior, dorsal/ventral and/or proximal/distal pattern specification; six HOXA (HOXA4, 5, 6, 7, 9, 10), five HOXB (HOXB2, 3, 5, 6, 8), two HOXC (HOXC4, 10), two HOXD (HOXD8, 11), as well as CDX1, LMX1B, LHX1, and BARX1 (Figure 1C, S2D). Seven motifs are from TFs important for limb specification, patterning or development, and seven motifs are from TFs involved in muscle development or differentiation (Figure 1C, S2D). MEOX2 is involved in both muscle and limb development when it acts in concert with PAX3 and SIX1/4 to activate *MYF5* in myoblasts migrating into the limb [28].

Having identified epigenetic differences in developmental TFs, we looked to see if the TFs are differentially expressed between the tissues. We performed RNA-seq on days 0 to 5 and day 12 of differentiation for the two quadricep and two TA cell lines (Fig S1A). We found 867 genes differentially expressed (|logFC| >1, FDR <0.01) including 54 TFs (Figure 1D). Of these TFs, 26 are related to axis patterning, 16 are involved in appendage development, and 12 are involved in muscle organ development (Figure S2E). The hindlimb determining TF *PITX1* is more highly expressed in TA than quadricep (Figure 1D) [29]. PITX1 activates *TBX4*, which was differentially methylated, in the developing hindlimb (Table S1) [8]. *TBX5*, which is responsible for specifying the forelimb, is more highly expressed in the quadricep than the TA (Figure 1D) [8]. Five of the differentially expressed TFs also had motifs which were significantly enriched in the differentially methylated regions; CDX1, MEOX2, HOXD8, FOXC1 and FOXD2 (Table S2). CDX1 is expressed in the limb bud and is responsive to retinoic acid (RA), WNT and FGF signals [30,31]. HOXD8 is expressed during early patterning in the proximal limb [8].

We compared the differences in DNA methylation with changes in gene expression by associating the differentially methylated regions with genes using GREAT (Methods) [32]. Seventeen of the TFs were both differentially expressed and differentially methylated (Figure 1E). This included *EN1*, which represses *WNT7A* in dorsal/ventral patterning, and *SALL4*, which regulates hindlimb outgrowth with TBX4 by regulating and *FGF10* [11,33]. Differences in methylation around *EN1* included proximal regions and the gene body that overlap annotated candidate cis-regulatory elements (Figure 1F). We compared the differentially expressed genes in TA and quadricep from our primary myoblasts from control and FSHD patients to those from RNA-seq from TA and quadricep patient biopsy samples [25]. *PITX1* and *IRX5* were upregulated in TA in both cell lines and biopsy samples for FSHD and control (Figure S2F). IRX5 expression in the hindlimb bud is important for specifying proximal and anterior regions of the limb [34]. Thus, muscle groups retain DNA methylation and expression differences in adult tissue based on early patterning.

### Genes induced upon myogenesis in FSHD are more highly expressed in more susceptible muscle

We previously identified a set of genes induced in FSHD2 cells upon differentiation up to day 5 (Williams & Jiang 2020). To assess FSHD specific gene upregulation in different muscle groups, we performed RNA-seq for both TA and quadricep from SMCHD1 mutated FSHD2. We found a set of 74 genes that are upregulated in both TA and quadricep FSHD2 cell lines starting around day 3 of differentiation (Figure 2A, 2B). This includes 47 of our original 54 FSHD-induced genes and an additional 27 genes (Figure 2B, S3A). Some of the new genes include previously identified DUX4 target genes such as *PRAMEF10* (Figure S2A) [21]. These additional genes appear to be more lowly expressed than the set we identified previously (Figure S2B). Interestingly, the quadricep, which is less susceptible to FSHD, has lower expression of these FSHD-induced genes (Figure 2A). However, when comparing FSHD biopsy samples from TA and quadricep, these genes are not significantly higher in the TA (Figure S3C, S3D).

**Figure 2:**
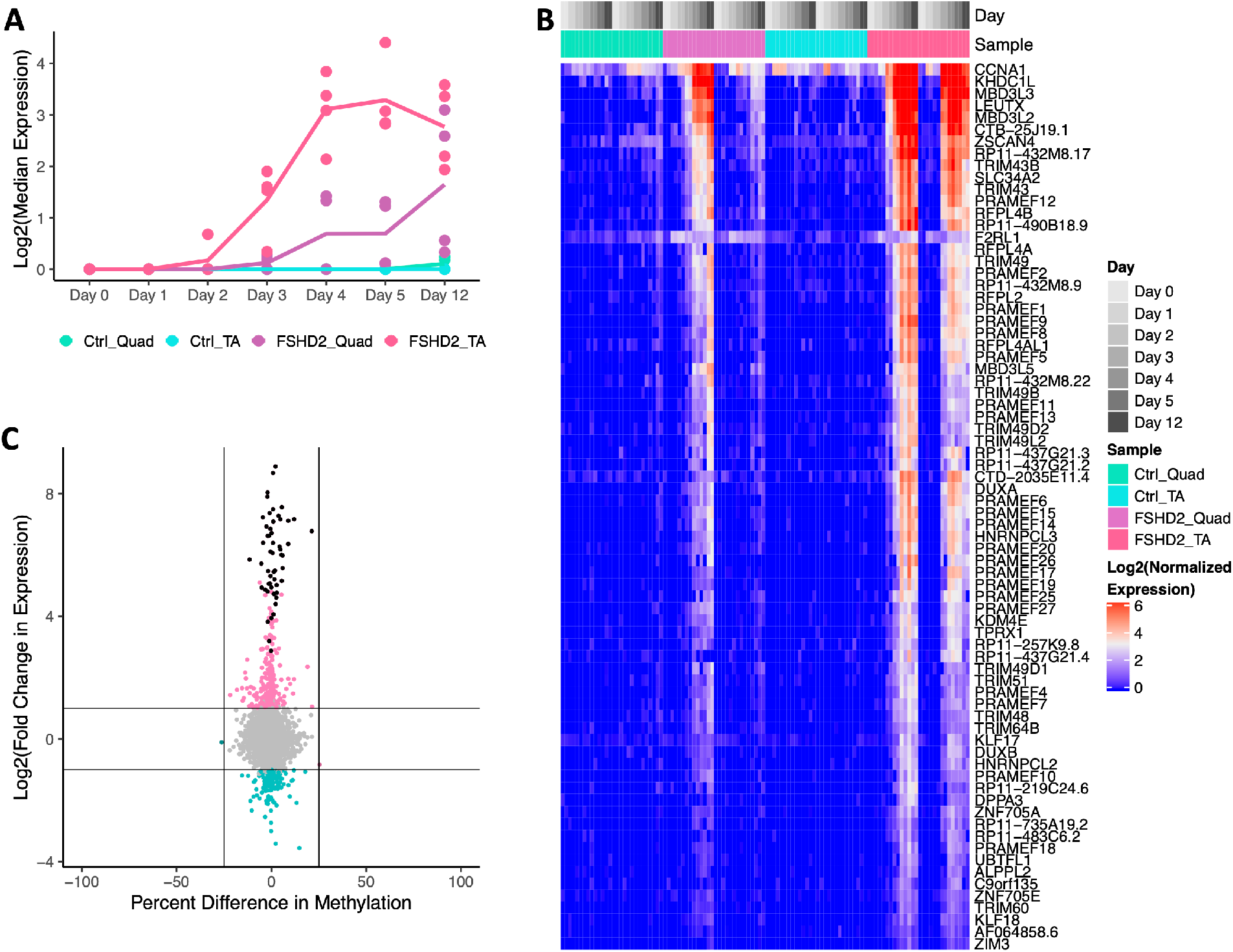
Promoters of FSHD-induced genes are not differentially methylated. **(A)** Median expression for given groups of 74 FSHD-induced genes with increased expression in FSHD2 during myogenesis. Dots represent median expression for individual samples. Line represents the mean for the four samples in each group. **(B)** Heatmap of the 74 FSHD-induced genes ordered by hierarchical clustering. **(C)** Scatterplot of differences in expression and promoter DNA methylation for FSHD2 versus control. Promoters are defined as -1.5kb to +0.5kb around the TSS. FSHD-induced genes are colored in black. Regions associated with genes with significant differences in gene expression but not DNA methylation are colored in pink and teal. Regions with significant differences in DNA methylation but not in expression of the associated gene are colored in dark teal and dark pink. Vertical lines intersect x-axis at -25 and 25 percent. Horizontal lines intersect y-axis at -1 and 1 log2(fold change).

We wanted to know whether differences in DNA methylation near these FSHD-induced genes could partially explain their strong upregulation. We summarized the promoter CpG sites for the 74 genes and recovered 53 gene promoters passing our coverage filters (Methods). The promoters of these genes are not significantly differentially methylated despite substantial increases in expression (Figure 2C). In fact, most of the promoters (33 out of 53) are highly methylated with greater than 50% methylation in both FSHD2 and control cell lines (Figure S3E). Hypermethylation in the promoter is generally associated with repressing gene expression, so we decided to determine whether DUX4 binds these promoter regions or could be regulating expression through binding of other regulatory regions. Eleven of the highly methylated FSHD-induced gene promoters overlap DUX4 binding sites determined by ChIP-seq (Figure S2D) [24].

To determine what TFs could be binding the methylated FSHD-induced gene promoters, we looked for enrichment of unmethylated and methylated TF motifs determined by SELEX [35]. While a methylated motif is not available for DUX4, the methylated motif for its paralog, DUXA, is highly enriched in 41 of the 74 promoters (Figure S3F, Table S3). Additionally, the methylated motif for OTX1, a PRD-like TF, is also enriched (Table S3). OTX1 has a similar motif to the DUX4 target TF LEUTX [36]. Performing enrichment using canonical motifs only, motifs for DUX4, DUXA, and OTX2, another PRD-like TF similar to LEUTX, are all enriched in the target promoters (Figure S3F, Table S3). The MEOX2 motif is also enriched, and *MEOX2* expression was significantly higher in quadricep than in TA (Figure 2A, Table S3).

### SMCHD1 mutation associated differences in DNA methylation and gene expression

To assess global methylation differences between SMCHD1 mutated FSHD2 and control cell lines, we compared all FSHD2 samples to all control samples and found 4527 regions more highly methylated in control and 3542 in FSHD2 (percent diff >10, qvalue <0.01) (Figure 3A). The top most differentially methylated region is *DBET*, which is close to *FRG2* and *DUX4* in the D4Z4 region on chromosome 4 that is hypomethylated in FSHD patients (Figure 3A) [37]. *DBET* is also less methylated in TA than quadricep in FSHD2 (Figure S4A). The D4Z4 region on chromosome 10 near *FRG2B* and *SYCE1* contains two hypomethylated regions in FSHD2, and *SYCE1* expression is upregulated in FSHD2 (Figure 3A, 3B) [38]. Hypomethylation in the D4Z4 region on chromosome 10 has been shown to be FSHD2 specific, and the upregulation of *SYCE1* that we observe is not seen in FSHD1 (Figure S4B) [39,40]. SMCHD1 has been shown to regulate clusters of genes such as the PCDH cluster on chromosome 5 (chr5:140,759,009-141,523,383, hg38), the SNRPN cluster (chr15:23,548,232-23,697,319, hg38), the rRNA cluster (chr1:228,552,374-228,653,525, hg38), and the tRNA cluster (chr1:161,395,860-161,624,746, hg38) [41,42]. Out of 142 regions in the PCDH cluster and 6 in the rRNA cluster, we find 27 regions and 3 regions, respectively, with hypomethylation in FSHD2 compared to control (Figure 3A). The SNRPN and tRNA clusters have negligible methylation differences (Figure 3A, Table S4). The hypomethylation observed in the PCDH and rRNA clusters is low (less than 25%) suggesting differences in methylation due to SMCHD1 heterozygosity are mild, especially when compared with methylation differences near D4Z4 on chromosomes 4 and 10.

**Figure 3:**
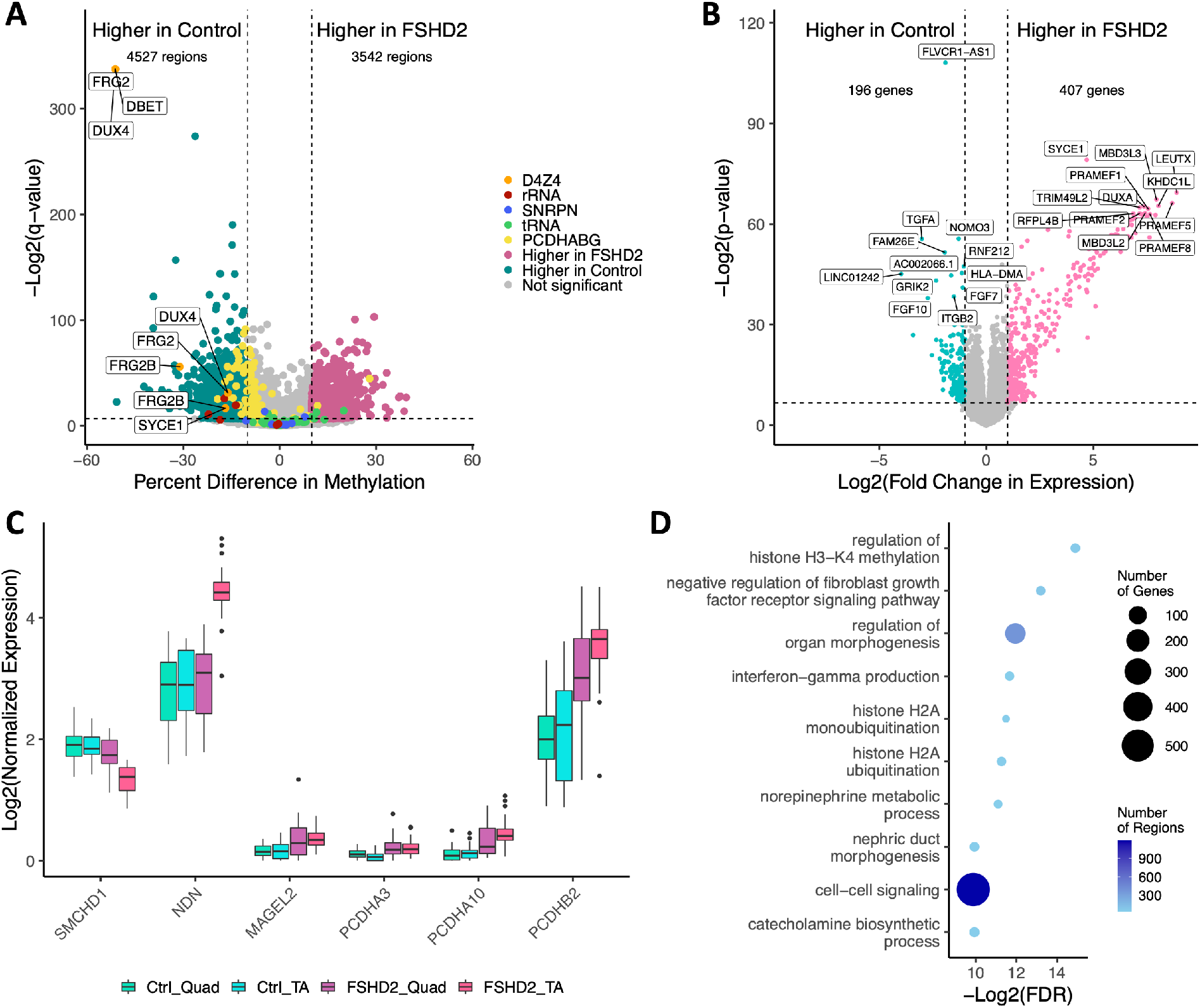
Global DNA methylation and gene expression differences in FSHD2 are associated with SMCHD1 mutations and signaling molecules. **(A)** Volcano plot of DNA methylation differences for FSHD2 versus control. Regions with a p-value of >0.01 or a percent methylation difference of <10 are taken as not significant and colored in grey. Regions with more methylation in FSHD2 are in dark pink. Regions with more methylation in quadricep are in dark teal. Regions within SMCHD1 regulated regions are colored according to the legend. Regions in the D4Z4 region on chromosome 4 or 12 have associated genes labelled. The numbers of regions higher in FSHD2 and control are labelled at the top. Vertical lines intersect y-axis at -10 and 10 percent. Horizontal line intersects y-axis at -log2(0.01). **(B)** Volcano plot of gene expression differences for FSHD2 versus control. Genes with a p-value of >0.01 or an absolute log2(fold change in expression) <1 are in grey. Genes with higher expression in the TA are in pink. Genes with higher expression in the quadricep are in teal. The numbers of genes with higher expression in FSHD2 or control are labelled at the top. The 12 genes with the lowest p-values for positive or negative fold change are labelled. Vertical lines intersect y-axis at -1 and 1 log2(fold change). Horizontal line intersects y-axis at -log2(0.01). **(C)** Boxplot of expression for genes in SMCHD1 regulated clusters and *SMCHD1*. **(D)** Gene ontology terms enriched in differentially methylated regions between FSHD2 and control. The top 10 significant terms (p-value <0.05) with at least 10 genes associated to the term from the background set of regions detected are shown. Color indicates number of differentially methylated regions in the term. Size indicates the number of genes associated with the differentially methylated regions in the term.

We looked at expression of genes in the PCDH and SNRPN clusters and found slight upregulation in FSHD2 samples (Figure 3C). Despite not having significant methylation differences, *NDN* and *MAGEL2* from the SNRPN cluster both have slightly elevated expression in FSHD2 (Figure 3C). Interestingly, *NDN* expression is much higher in FSHD2 cells from the more susceptible muscle group TA (Figure 3C). Three genes from the PCDH cluster were upregulated in FSHD2 cells consistent with the hypomethylation in this region (Figure 3C). *SMCHD1* expression is slightly lower in FSHD2 samples than control, which is surprising since *SMCHD1* mutations in these FSHD2 patients affect protein function, not expression (Figure 3C).

We assessed global expression differences between FSHD2 and control combining all differentiation days. The most highly upregulated genes in FSHD2 include the FSHD-induced genes previously identified (Figure 3B). The most significantly downregulated gene in FSHD2 is *FLVCR1-AS1*, a lncRNA, which can act as a sponge for miRNAs that inhibit proliferation (Figure 3B) [43]. The genes for signaling molecules *FGF7, FGF10* and *TGFA* are also down regulated in FSHD2 compared to control (Figure 3B, Table S5). Using gene ontology analysis, the differentially methylated regions were identified as being associated with genes involved in histone modifications, including H3K4 methylation and H2A ubiquitination, and FGF signaling (Figure 3D).

### LTR loci are highly expressed in muscle that is more susceptible to FSHD

DUX4 activates expression of transposable elements (TEs), and we therefore wanted to assess the extent of TE upregulation in our samples [24,44]. In FSHD2 cells, 580 repeat loci comprised of 175 different TE types are upregulated in a time specific manner starting at day 3 of differentiation (Figure 4A, 4B). Of these 175 types, 79 are LTRs including significant enrichment of ERVL-MaLR (p=4.9e-92), ERVL (p=2.6e-17) and ERV1 (p=2.7e-5) (Figure 4C). Specific repeat types, such as THE1D, MLT2A1 and THE1C, which are regulated by DUX4, are also significantly enriched and upregulated (Figure S5A) [24]. These loci are substantially upregulated in FSHD2 TA and show only slight upregulation in quadricep at day 12 of differentiation (Figure 4A).

**Figure 4:**
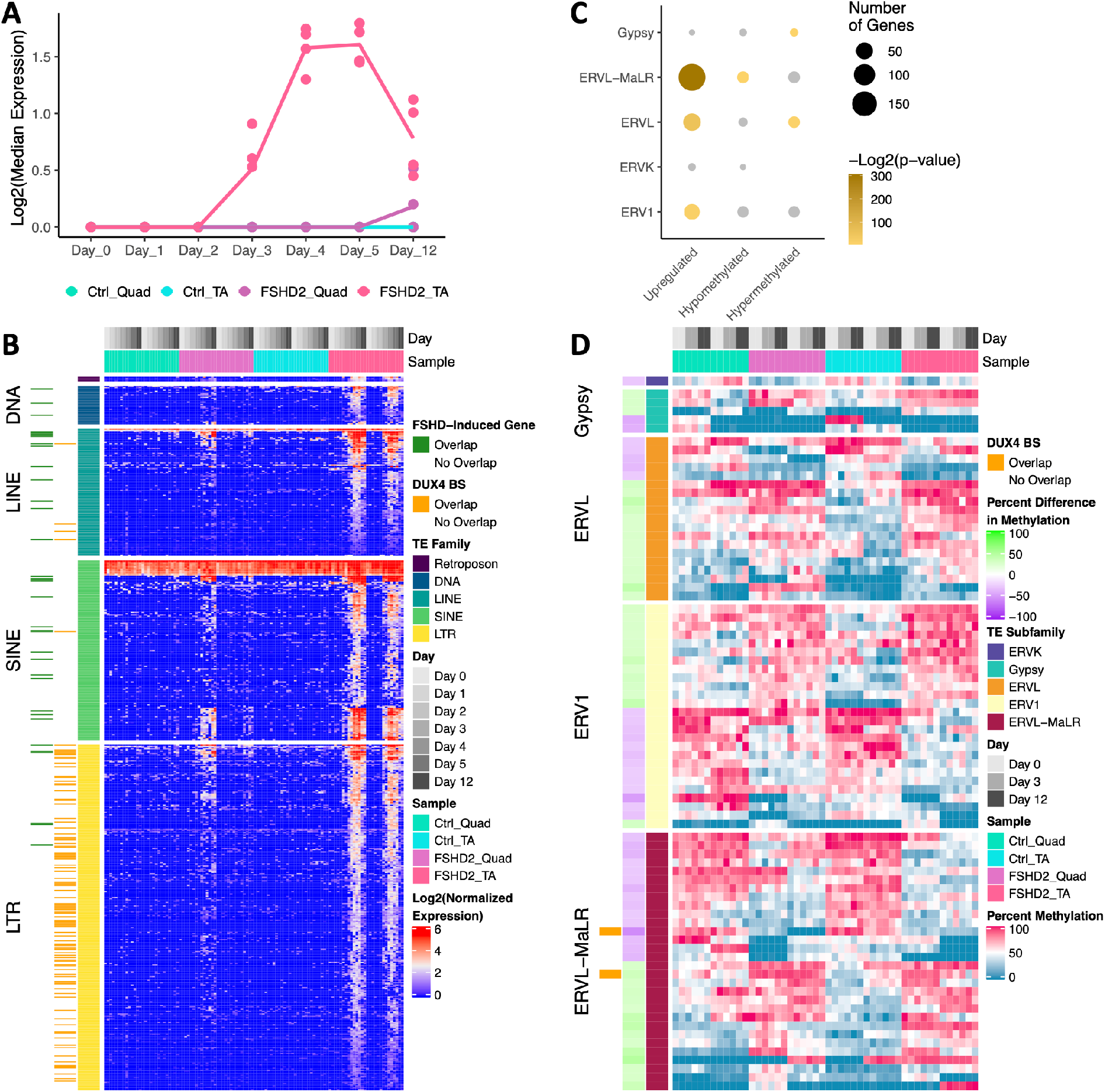
LTRs upregulated and hypomethylated in FSHD2. **(A)** Median expression for given groups of 580 TEs with increased expression in FSHD2 during myogenesis. Dots represent median expression for individual samples. Line represents the mean for the four samples in each group. **(B)** Heatmap of expression of TEs with increased expression in FSHD2 during myogenesis split by class of TE. Loci that overlap FSHD-induced genes are indicated in green on the left. Loci which overlap known DUX4 binding sites are labelled in orange. **(C)** Enrichment of subclasses of LTRs in TEs upregulated in FSHD2 or differentially methylated in FSHD2. Classes with p-value >0.05 are in grey. Size indicates number of loci in category. **(D)** Heatmap of percent methylation for LTR loci with significant methylation differences between FSHD2 and control split by LTR subclass. Loci which overlap a known DUX4 binding site are indicated in orange on the left.

These upregulated TEs significantly overlap (p=2.3e-70) DUX4 binding sites identified by ChIP-seq (Figure 4B) [24]. Out of 104 loci that overlap DUX4 binding sites, 99 are LTRs (p=2.1e-28) including 79 ERVL-MaLRs (p=5.7e-24) (Figure 4B, 4C). To determine whether the TEs are upregulated because they overlap FSHD-induced genes, we intersected the TE loci with the FSHD-induced genes. Thirty-six TE loci overlap the FSHD-induced genes, only 6 of which are LTRs (Figure 4B). A previous study found that LTRs bound by DUX4 could act as alternative promoters for protein coding and lncRNAs [24]. We find that 20 out of the 79 LTRs upregulated in FSHD2 overlap LTRs identified as alternative promoters, including 15 for protein coding genes such as *MLT1E1A-NT5C1B* and 5 for lncRNAs (Table S6).

We summarized CpG sites over LTR loci to determine whether DUX4-regulated LTRs are differentially methylated (Methods). Between FSHD2 and control, 45 LTRs are more highly methylated (percent diff >25%, qvalue <0.01) in FSHD2, and 36 are more highly methylated in control (Figure 4D). These loci were not in the set identified as upregulated in FSHD2. Loci with more methylation in FSHD2 are enriched for ERVL (p=0.01) and Gypsy (p=0.04) elements (Figure 4C). FSHD2 hypomethylated TEs are enriched for ERVL-MaLR (p=0.03) which are enriched in the set of TEs upregulated in FSHD2 (Figure 4C). Additionally, two ERVL-MaLR loci overlap DUX4 binding sites (Figure 4D).

### Muscle group specific gene expression in FSHD

To identify transcriptional differences between muscle groups that may cause or result from differential susceptibility to FSHD, we performed RNA-seq on FSHD1 patient matched deltoid and bicep derived cell lines for days 0 and 5 of differentiation as well as from TA for day 0 (Figure S1). We identified muscle specific expression patterns by performing pairwise comparisons between the muscle groups at day 0 and taking the intersect of genes specific for one muscle group (Methods). A total of 582 unique genes were specific to a muscle group with either especially high or low expression for the group (Figure 5A). Bicep specifically expresses *LHX8* which can regulate *SHH* expression in development of various tissues in the upper body (Figure 5B) [45]. Deltoid expresses *HOXD3* which is involved in limb outgrowth (Figure 5B) [46]. TA has higher expression of *EYA1* which is involved in activating PAX3 and MYF5 in limb muscles (Figure 5B) [5]. Quadricep has high expression of *TLL1* which is involved in dorsal ventral patterning and can indirectly regulate myostatin (Figure 5B) [47,48]. Interestingly, we see the hindlimb specific TF *TBX5* is high in the bicep, deltoid and quadricep but not the tibialis anterior (Figure S6A).

**Figure 5:**
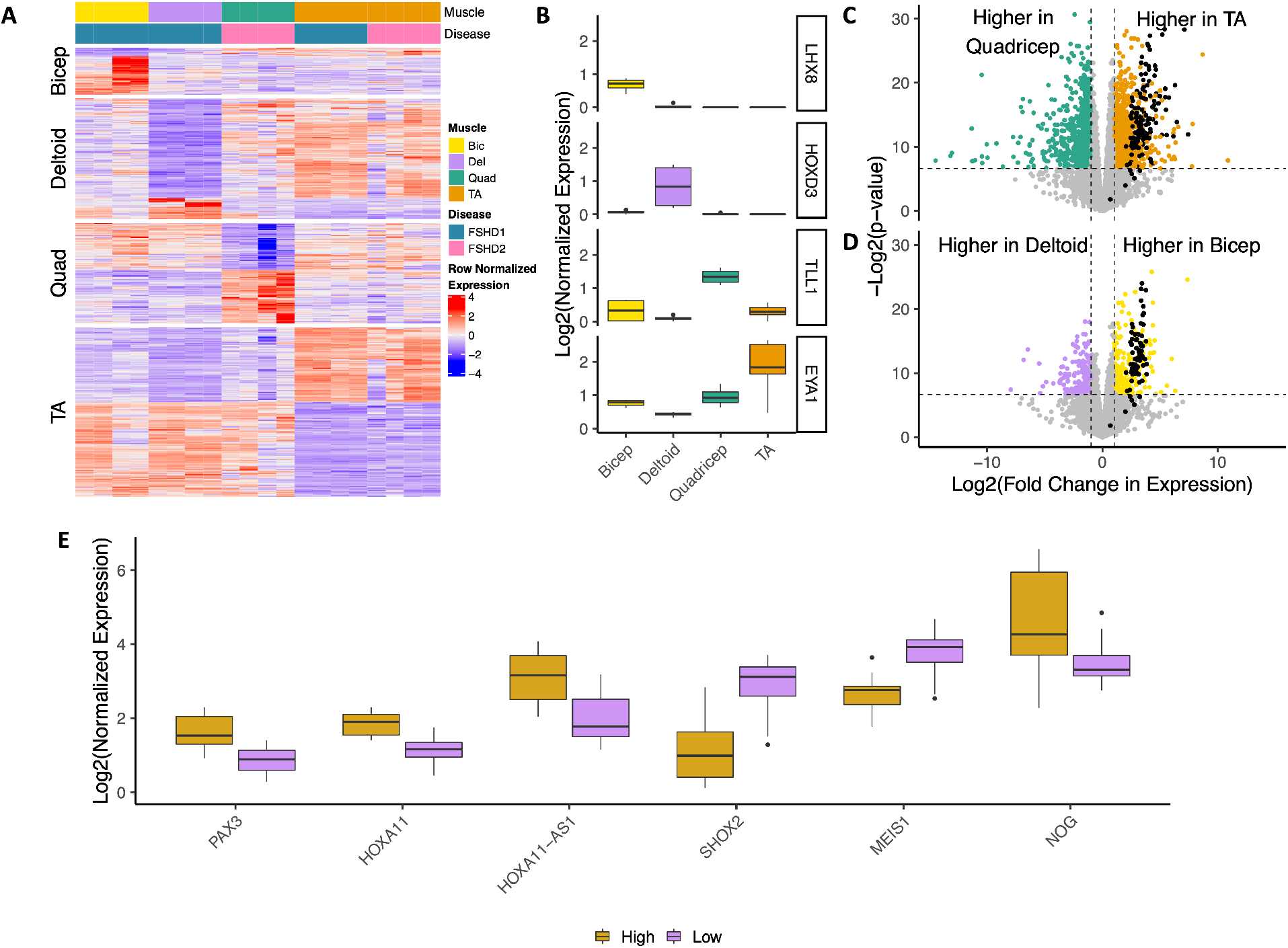
Muscle group specific expression in muscles more and less susceptible to FSHD. **(A)** Heatmap of genes with muscle group specific expression. Top color bars indicate FSHD1 or FSHD2 and the muscle group of origin. **(B)** Boxplot of expression of four genes with muscle group specific expression. **(C)** Volcano plot of gene expression differences for FSHD2 TA versus FSHD2 quadricep at day 5 of differentiation. FSHD-induced genes are colored in black. Genes with a p-value of >0.01 or an absolute log2(fold change in expression) <1 are in grey. Genes with higher expression in the TA or quadricep are in orange or green, respectively. Vertical lines intersect y-axis at -1 and 1 log2(fold change). Horizontal line intersects y-axis at - log2(0.01). **(D)** Volcano plot of gene expression differences for FSHD1 bicep versus FSHD1 deltoid at day 5 of differentiation. FSHD-induced genes are colored in black. Genes with a p-value of >0.01 or an absolute log2(fold change in expression) <1 are in grey. Genes with higher expression in the bicep or deltoid are in yellow or purple, respectively. Vertical lines intersect y-axis at -1 and 1 log2(fold change). Horizontal line intersects y-axis at -log2(0.01). **(E)** Boxplot of genes with differences in expression in muscle groups with high or low susceptibility to FSHD.

FSHD tends to affect upper musculature first, so we compared the two forelimb muscles (bicep and deltoid) to the two hindlimb ones (quadricep and TA). Genes in the 5’ end of the HOXC cluster, including *HOXC9* through *13* as well as *HOXC-AS1, HOXC-AS5* and *HOTAIR*, were all more highly expressed on average in the hindlimb than the forelimb (Figure S6B, S6C). Genes in the 3’ end of the HOXD cluster are more highly expressed in the forelimb (Figure S6C).

We then compared expression between the commonly and less affected muscle groups in the forelimb and hindlimb separately to investigate differences between muscle groups more susceptible to FSHD. As noted previously, FSHD-induced genes are more highly expressed in TA than quadricep, 71 out of 74 at day 5 of differentiation (Figure 5C). FSHD-induced genes are also more highly expressed in bicep, which is more commonly affected, than deltoid (67 out of 74) (Figure 5D). Higher expression of DUX4 target genes has been shown previously to correlate with active signs of disease [25]. Here we show that the FSHD gene signature is also more highly expressed in more susceptible tissue.

To find genes specific to commonly affected groups, we took the intersect of the genes differentially expressed in bicep compared to deltoid and TA compared to quadricep in the same direction (Methods). This yields 28 genes more highly expressed in commonly affected groups, and 27 higher in the less affected groups including 7 TFs total (Figure S6D, Table S7). The myogenic precursor TF *PAX3* and *HOXA11* are more highly expressed in more susceptible groups (Figure 5E). TFs more highly expressed in the less susceptible muscles include *SHOX2* and *MEIS1. MEIS1* and *SHOX2* are expressed in the proximal region of the limb [8,9,49]. *PAX3* marks myogenic precursor cells and may play a role in satellite cells postnatally [5]. *HOXA11* is expressed in the zeugopod in development [8,10]. We also see higher expression of genes involved in WNT signaling including *Noggin* (Figure 5E, Figure S6C). Also higher in more affected tissues is *HEYL* which is involved in NOTCH signaling, represses MYOD expression, and is required for satellite cell proliferation in a model of hypertrophy (Figure S6C) [50,51]. In conclusion, we identified DNA methylation and transcriptional differences between muscle groups in cell lines derived from adult tissue.

## Discussion

We identified developmental transcription factors with differences in DNA methylation and expression in cells derived from adult tissue. By examining cells from FSHD patients, we found differences between muscle groups that are related to disease, including *PITX1*, emphasizing the consideration of sample origin in transcriptome studies in FSHD. Genes induced specifically in FSHD are not differentially methylated, but genes regulated by SMCHD1 are hypomethylated and slightly overexpressed in SMCHD1 mutated FSHD2 patient cells. Importantly, we find a set of genes that possibly correlate with muscle group susceptibility to FSHD.

The differential DNA methylation and gene expression between muscle groups supports a potential role for developmental transcription factors in adult tissue. Previous work in mice showed transcriptional and methylation differences between extraocular muscles (EOM) and TA and that transplantation could alter the location-specific transcriptional profiles [15]. Notably, EOM cells, which do not express HOX genes, upregulate TA specific HOX genes upon transplantation supporting a role for environmental stimuli in controlling location-associated TFs [15]. Notably, *PITX2* expression in EOM cells was resistant to change after transplantation to the TA, and PITX2 can increase the ability of satellite cells to regenerate [4,15]. We find substantial methylation differences in PITX2 between TA and quadricep. The role of some of these other TFs in adult tissue warrants further investigation.

The transcriptional differences that we observe between the bicep and TA compared with the deltoid and quadricep may be the result of inherent differences between these muscle groups or due specifically to disease. While the differences that we observed between TA and quadricep could be due to individual differences, our bicep and deltoid samples were taken from the same individuals. Comparison of these muscle groups from healthy individuals could identify whether differences between the susceptible groups are due to inherent differences, active disease or differences inherent to FSHD patients. The higher expression of FSHD-induced genes in the more commonly affected muscle groups may be due to either active disease or inherent differences between the groups. Indeed, FSHD-related gene expression has been shown to be higher in tissue with signs of active disease [25,26]. Differences in expression of homeobox transcription factors between commonly and less affected groups suggests a possible link to the governing of muscle groups affected.

A previous study of DNA methylation in FSHD and control patients found relatively little differential methylation compared to what we observe, which may be the result of increased sample size or to the SMCHD1 mutations in the FSHD2 patients [52]. The enrichment in hypomethylation of ERVL-MaLRs suggests a mechanism for their increased expression in FSHD. The differentially methylated ERVL-MaLRs do not significantly overlap DUX4 binding sites, which suggests that regulation of DNA methylation in those regions is independent of DUX4 binding but could possibly be attributed to SMCHD1 mutations. In addition, we noted differences in methylation related to FGF signaling and downregulation of two *FGFs* and *TGFα* in FSHD2 cells. FGF10 is an important contributor to limb outgrowth but is largely unexplored in FSHD. FGF1 and FGF2 were found in a biopsy of one patient with a severe phenotype [53]. We found *NOG* more highly expressed in more susceptible muscle groups in FSHD. Noggin inhibits multiple BMPs including BMP4 [54]. BMP4 can activate *FGF7* and *FGF10* expression [55].

The hypermethylation and lack of difference in the methylation of the FSHD-induced gene promoters could indicate several possibilities. First, DUX4 most likely binds other regulatory elements outside of these promoter regions, such as enhancers, such that highly methylated promoters are not prohibitive to activation by DUX4. Second, DUX4 and/or other binding partners may not be sensitive to DNA methylation. Indeed, homeobox transcription factors preferentially bind methylated DNA, and we find the methylated motif for DUXA enriched at the promoters of FSHD-induced genes [35]. Second, complexes that are indirectly called to methylated DNA, such as SIN3, regulate *DUX4* expression, and their targets are affected in DUX4-affected cells [56–59]. DUX4 also upregulates the *MBD3L* genes which are methyl binding domain proteins, and MBD3L2 is known to bind methylated DNA [60,61]. Third, demethylated promoters may only be present in the subset of nuclei that activate the DUX4 program. As affected FSHD nuclei make up a small percentage of total nuclei, differences in those nuclei would only be observable with single nucleus methylation assays [23,57].

In summary, we identified transcriptional and DNA methylation differences between muscle groups from adult tissue of healthy individuals and FSHD patients. We identified a set of genes potentially linked to susceptibility or progression of affected muscle groups in FSHD, including genes specifically upregulated in FSHD across myogenesis. Understanding the genomic basis of susceptibility is important for identifying key mechanisms of FSHD progression.

## Methods

### Human myoblast culture and differentiation

Human control, FSHD1 and FSHD2 myoblast cells from patient quadricep, tibialis anterior, bicep and deltoid biopsies were grown as previously described [57]. Cells were cultured on dishes coated with collagen in high glucose DMEM (Gibco) supplemented with 20% FBS (Omega Scientific, Inc.), 1% Pen-Strep (Gibco), and 2% Ultrasor G (Crescent Chemical Co.). Day 0 cells were kept at low confluency to prevent spontaneous differentiation. Upon reaching 80% confluence, differentiation was induced by using high glucose DMEM medium supplemented with 2% FBS and ITS supplement (insulin 0.1%, 0.000067% sodium selenite, 0.055% transferrin; Invitrogen). Fresh differentiation medium was changed every 24 hours.

### DNA methylation library preparation

Genomic DNA was collected from a single 6 cm dish using the DNEasy Blood & Tissue kit (69504, Qiagen). For both reps of day 0, day 3 and rep 1 of day 12 for Control-4, gDNA from two plates were pooled and concentrated using DNA Clean & Concentrator-25 (D4033, Zymo) to obtain high enough input and concentration. All DNA methylation data was generated using the TruSeq Methyl Capture EPIC kit (FC-151-1003, Illumina) according to manufacturer’s protocol [62]. Libraries were sequenced on the Illumina NextSeq500 with paired-end 75 bp reads to a depth of 20 to 82 million reads.

### DNA methylation data processing

Raw reads from TruSeq Methyl Capture EPIC libraries were mapped to canonical chromosomes from hg38 and the patch region of D4Z4 (chr4_KQ983257v1_fix) using bismark (version 0.19.0) [63]. Sites were extracted from the bam file using bismark_methylation_extractor with paired end and no overlap specified. To remove bias from the ends of reads, 2 bp from the 5’ end of read 2 and 1 bp from the 3’ end of read 1 were ignored when extracting sites. All CpG sites were read into methylKit (version 1.16.0) [64]. Sites within 200 bp were merged into one region using bumphunter (version 1.32.0) [65]. Methylation over those regions was summarized using regionCounts and filtered for at least three CpG sites. Regions were then filtered for a coverage of at least 5. Regions were normalized using normalizeCoverage, and only regions with coverage in all samples were kept. Differential methylation was calculated using calculateDiffMeth with a chi-squared test with basic overdispersion correction. Differentially methylated regions (DMRs) were defined with a q-value <0.01 and a percent difference in methylation of at least 25% for TA versus quadricep and FSHD2 versus control for TE loci but 10% for FSHD2 versus control.

Regions were associated with genes and gene ontology using rGREAT (version 1.20.0) [32] with submitGreatJob with hg38 specified. GO term plots include the top 10 terms with greater than 10 genes associated with the background set for the term and a Benjamini-Hochberg corrected p-value of less than 0.05. Regions were overlapped with SMCHD1 regulated regions and DUX4 binding sites from [24] using findOverlaps from GenomicRanges (version 1.40.0) [66].

For the promoter analysis, CpG sites were summarized for 1.5 kb upstream and 0.5 kb downstream of the TSS using regionCounts and filtered for at least 1 CpG site. Promoters were filtered for a coverage of at least 5 then normalized. Promoters detected in all samples were kept for further analysis as described.

### Motif analysis

Motif enrichment was performed on DMRs extended 24 bp in both directions since the regions end on CpG sites. Enrichment was performed using AME (meme, version 5.2.0) [67] on JASPAR 2020 core redundant motifs from human only [68]. Association of TFs with GO terms for figure 1C was done by associating the TFs with the following GO terms: limb morphogenesis, embryonic limb morphogenesis, embryonic forelimb morphogenesis, limb development, limb bud formation for limb; anterior/posterior pattern specification, proximal/distal pattern formation, dorsal/ventral pattern formation, dorsal/ventral pattern specification for axis patterning; skeletal muscle tissue development, skeletal system development, negative regulation of myoblast differentiation, muscle organ development, skeletal muscle cell differentiation, skeletal muscle tissue regeneration, myoblast development, and positive regulation of myoblast proliferation for muscle. Upset plots were created using UpsetR (version 1.4.0) [69].

### TE methylation analysis

For the TE analysis, CpG sites were summarized over LTR loci from repeatmasker taken from UCSC and filtered for at least 1 CpG site. Loci were filtered for a coverage of at least 5 then normalized. Loci detected in all samples were kept for further analysis as described.

### RNA sequencing library preparation

Total RNA was collected using the RNEasy kit (74106, Qiagen). RNA was converted to cDNA using the Smart-Seq2 protocol [70]. Libraries were constructed with the Nextera DNA Library Prep Kit (Illumina) for days 0 to 5 for Control-3, Control-4, FSHD2-2 and FSHD2-3. The Nextera DNA Flex Library Prep Kit (Illumina) was used for all other RNA-seq samples. Libraries were sequenced on the Illumina NextSeq500 with paired-end 43 bp reads to a depth of 5 to 40 million reads. Data for days 0 through 5 for Control-1, Control-2, Control-3, Control-4, FSHD2-1 and FSHD2-2 were obtained from GEO (accession number GSE143493).

### RNA sequencing data processing

Raw reads from bulk RNA-seq were mapped to hg38 by STAR (version 2.5.1b) [71] using defaults except with a maximum of 10 mismatches per pair, a ratio of mismatches to read length of 0.07, and a maximum of 10 multiple alignments. Quantitation was performed using RSEM (version 1.2.31) [72] with defaults with gene annotations for protein coding genes and lncRNAs from GENCODE v28. Counts were batch corrected for the two different library prep methods using ComBat-seq from sva (version 3.36.0) [73]. Genes were filtered for 10 counts in all samples of at least one condition (same FSHD status, muscle group and differentiation day) using filterByExpr from edgeR (version 3.30.3) [74]. Normalization for days 0 to 5 and 12 for control and FSHD2 for figures 1 through 3 was performed separately from days 0 and 5 for FSHD2 and FSHD1 for figure 5. TMM normalized counts from edgeR were used for differential expression analysis in edgeR and clustering genes into similar expression profiles using maSigPro (version 1.60.0) [75]. TMM normalized counts were TPM normalized for plotting using effective gene lengths for each sample calculated by RSEM. Heatmaps were created using ComplexHeatmap (version 2.4.3) [76]. Median expression of gene groups for each sample was calculated by taking the median TPM normalized TMM values for all genes, and lines represent the mean of the median values for the given samples. Muscle specific genes were identified by performing pairwise comparisons between muscle groups (i.e. TA vs quadricep, TA vs deltoid, TA vs bicep) as described with edgeR and taking the intersect of all genes specific to the same muscle group in all comparisons (i.e. up in all TA comparisons). Genes with higher expression in more or less susceptible tissue were identified by intersecting the genes higher in TA than quadricep and higher in bicep than deltoid and vice versa. The list of transcription factors was taken from AnimalTFDB (version 3.0) [77].

### TE mapping and processing

To map to TEs, fastqs were aligned as described above to the full GENCODE v28 annotation with a maximum of 100 multiple alignments and maximum of 100 loci anchors. Reads mapping to TE loci from repeatmasker from UCSC were estimated using featureCounts (subread version 2.0.1) [78] with fractional counts for multimapped reads for paired end reads. Genes were filtered for 2 counts in all samples of at least one condition (same FSHD status, muscle group and differentiation day) using filterByExpr from edgeR (version 3.30.3) [74]. Counts were normalized as described above. Loci were clustered into similar expression profiles as described above. Loci were overlapped with FSHD-induced genes (TSS to TES) or DUX4 binding sites from [24] using findOverlaps from GenomicRanges (version 1.40.0) [66]. Fisher’s exact test (stats version 4.0.2) was used for enrichment of TE classes and DUX4 binding site overlaps with the “greater” alternative hypothesis.

### Processing of publicly available data

Fastqs for the biopsy RNA-seq data from [21,25,79,80] were obtained from GEO with the accessions in table 1. Fastqs were processed in the same way as bulk RNA-seq. Counts were normalized as described above but with no batch correction.

**Table 1:** GEO Accession numbers for fastqs from previous studies.

| Disease | Muscle | SRA | Reference |
| --- | --- | --- | --- |
| Ctrl | Quad | SRR7293780 | [25] |
| Ctrl | Quad | SRR7293781 | [25] |
| Ctrl | Quad | SRR7293782 | [25] |
| Ctrl | Quad | SRR7293783 | [25] |
| Ctrl | Quad | SRR7293784 | [25] |
| Ctrl | Quad | SRR7293785 | [25] |
| Ctrl | Quad | SRR7293793 | [25] |
| Ctrl | Quad | SRR7293794 | [25] |
| Ctrl | Quad | SRR7293795 | [25] |
| FSHD | Quad | SRR7293761 | [25] |
| FSHD | Quad | SRR7293769 | [25] |
| FSHD | Quad | SRR7293774 | [25] |
| FSHD | Quad | SRR7293786 | [25] |
| FSHD | Quad | SRR7293788 | [25] |
| FSHD | Quad | SRR7293801 | [25] |
| FSHD | TA | SRR7293759 | [25] |
| FSHD | TA | SRR7293763 | [25] |
| FSHD | TA | SRR7293766 | [25] |
| FSHD | TA | SRR7293767 | [25] |
| FSHD | TA | SRR7293770 | [25] |
| FSHD | TA | SRR7293771 | [25] |
| FSHD | TA | SRR7293787 | [25] |
| FSHD | TA | SRR7293790 | [25] |
| FSHD | TA | SRR7293791 | [25] |
| FSHD | TA | SRR7293792 | [25] |
| FSHD | TA | SRR7293796 | [25] |
| FSHD | TA | SRR7293799 | [25] |
| FSHD | TA | SRR7293800 | [25] |
| MB135_HDUX4CA_nodox_rep1 | NA | SRR4019004 | [79] |
| MB135_HDUX4CA_WITHdox_rep1 | NA | SRR4019005 | [79] |
| MB135_HDUX4CA_nodox_rep2 | NA | SRR4019006 | [79] |
| MB135_HDUX4CA_WITHdox_rep2 | NA | SRR4019007 | [79] |
| MB135_HDUX4CA_nodox_rep3 | NA | SRR4019008 | [79] |
| MB135_HDUX4CA_WITHdox_rep3 | NA | SRR4019009 | [79] |
| FSHD_1_1_neg | NA | SRR2020583 | [80] |
| FSHD_1_2_neg | NA | SRR2020584 | [80] |
| FSHD_2_2_BFP | NA | SRR2020585 | [80] |
| FSHD_2_3_BFP | NA | SRR2020586 | [80] |
| FSHD_1_3_neg | NA | SRR2020587 | [80] |
| FSHD_1_1_BFP | NA | SRR2020588 | [80] |
| FSHD_1_2_BFP | NA | SRR2020589 | [80] |
| FSHD_1_3_BFP | NA | SRR2020590 | [80] |
| FSHD_2_1_neg | NA | SRR2020591 | [80] |
| FSHD_2_2_neg | NA | SRR2020592 | [80] |
| FSHD_2_3_neg | NA | SRR2020593 | [80] |
| FSHD_2_1_BFP | NA | SRR2020594 | [80] |
| Control_20_Mt | NA | SRR1398556 | [21] |
| Control_21_Mb | NA | SRR1398557 | [21] |
| Control_21_Mt | NA | SRR1398558 | [21] |
| Control_22_Mb | NA | SRR1398559 | [21] |
| Control_22_Mt | NA | SRR1398560 | [21] |
| FSHD2_12_Mt | NA | SRR1398561 | [21] |
| FSHD2_14_Mb | NA | SRR1398562 | [21] |
| FSHD2_14_Mt | NA | SRR1398563 | [21] |
| FSHD2_20_Mb | NA | SRR1398564 | [21] |
| FSHD2_20_Mt | NA | SRR1398565 | [21] |
| FSHD1_4_Mb | NA | SRR1398566 | [21] |
| FSHD1_4_Mt | NA | SRR1398567 | [21] |
| FSHD1_6_Mb | NA | SRR1398568 | [21] |
| FSHD1_6_Mt | NA | SRR1398569 | [21] |

## Data Availability

Data generated for this manuscript are available through GEO (GSE174370) or through dbGap (accession number TBD).

## Authors’ contributions

KW – Conceptualization, Data curation, Formal analysis, Investigation, Writing – original draft, Writing – review & editing

XK – Investigation, Resources

NVN – Investigation

CM – Resources

RT – Resources

KY – Funding acquisition, Project administration, Resources, Supervision, Writing – review & editing

AM – Funding acquisition, Project administration, Resources, Supervision, Writing – review & editing

## Acknowledgements

We would like to thank the NIH UMMS Senator Paul D. Wellstone Muscular Dystrophy Cooperative Research Center for FSHD Research for providing the bicep and deltoid cell lines.

---

**Figure S1:**
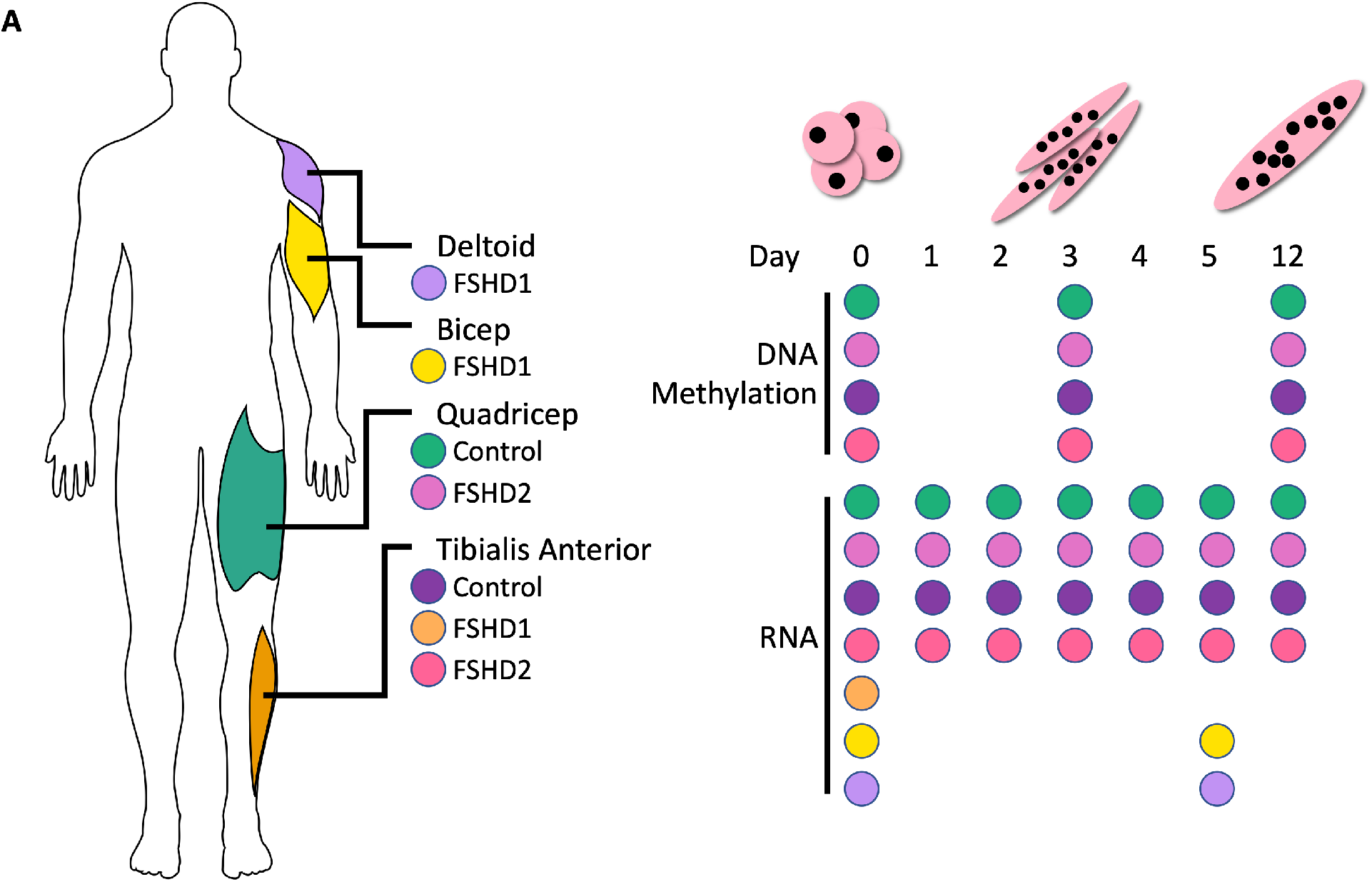
Experiment overview. **(A)** Overview of samples used for bisulfite and RNA sequencing.

**Figure S2:**
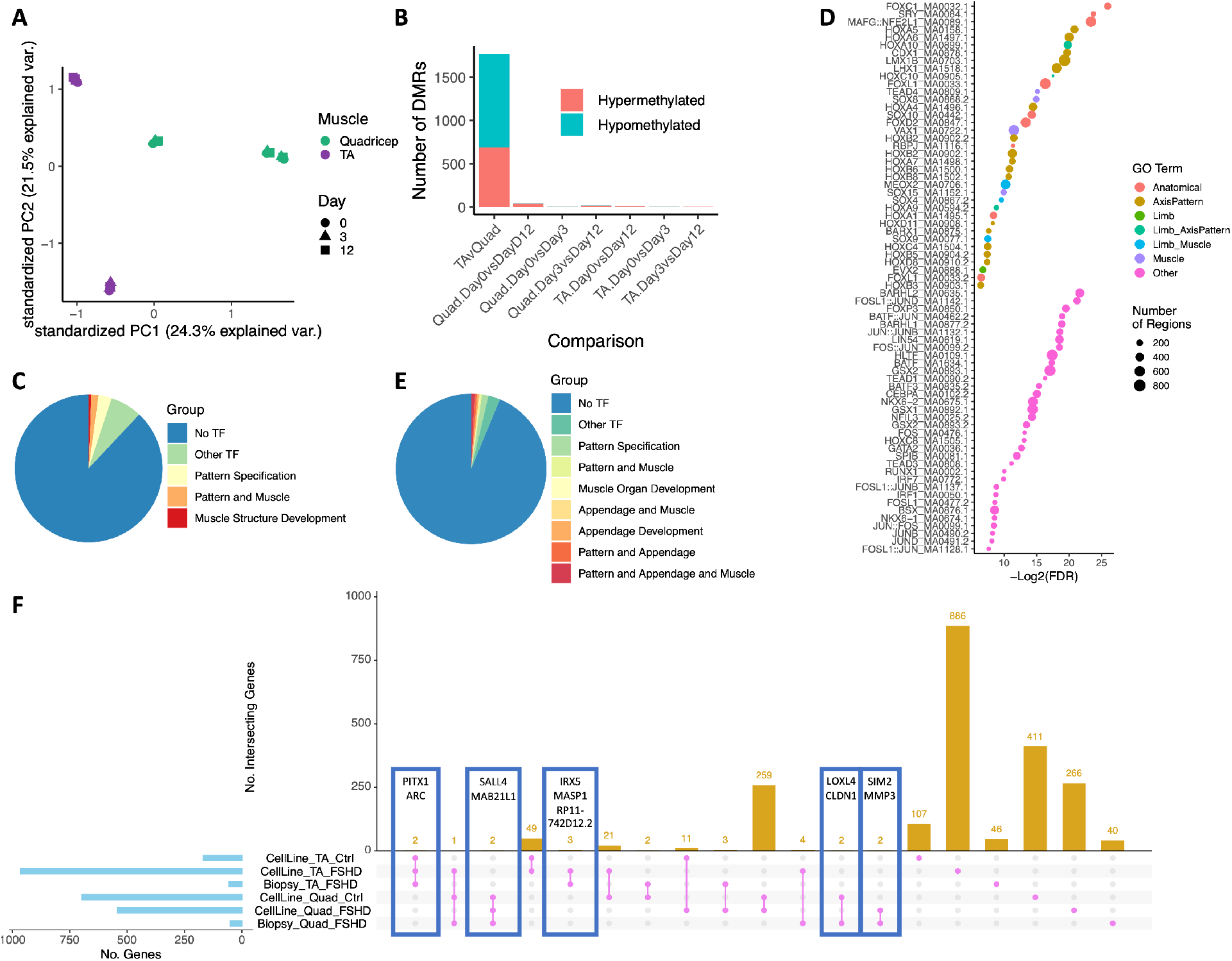
DNA methylation and gene expression differences for TA compared to quadricep. **(A)** Principal component analysis (PCA) of control TA and quadricep DNA methylation. **(B)** Stacked barplot of the number of differentially methylated regions between TA and quadricep. **(C)** Pie chart of the number of differentially methylated regions associated with genes for transcription factors. Regions not associated with a transcription factor are in blue. Regions associated with transcription factors are colored by annotated gene ontology terms for pattern specification or muscle structure development. **(D)** Transcription factor motifs enriched in differentially methylated regions between TA and quadricep. Color indicates the gene ontology annotation associated with transcription factor, either anatomical development, axis patterning, limb development, or muscle development (see Methods for specific GO terms). Size indicates number of regions with given motif. FDR cutoff of 0.05. **(E)** Pie chart of the number of differentially expressed transcription factors for TA versus quadricep. Genes that are not transcription factors are in blue. Transcription factors are colored by annotated gene ontology terms for pattern specification, appendage development or muscle structure development. **(F)** Upset plot of genes differentially expressed between TA and quadricep in control cells, FSHD2 cells or in FSHD biopsies from [26]. Number of genes higher in the given muscle in the given comparison are indicated in the light blue barplot on the left. Number of genes found in the intersections indicated in magenta are given in the gold barplot at top. Genes in intersections outlined in blue are labelled.

**Figure S3:**
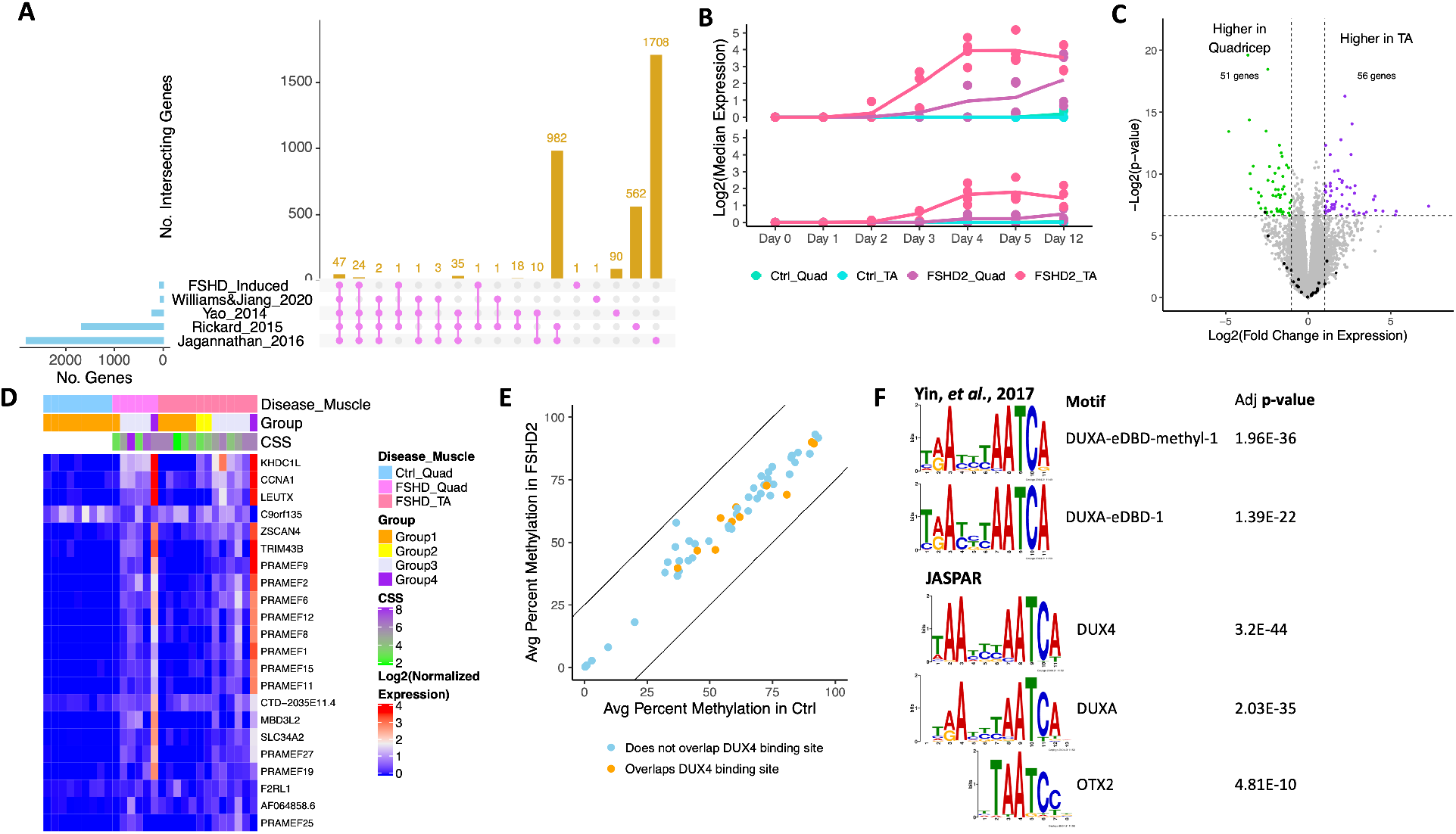
Assessment of FSHD-induced genes. **(A)** Upset plot of FSHD-induced genes compared with the gene list from [53] and reanalyzed data from [21,75,76]. Yao 2014 is a comparison of differentiated FSHD and control myotubes [21]. Rickard 2015 is a comparison of DUX4 expressing myocytes identified by a reporter to those negative for the reporter [76]. Jagannathan 2016 is a comparison of induced expression of DUX4 to non-induced [75]. **(B)** Median expression for given groups of 74 FSHD-induced genes. Those identified in [53] are on top, and those new from this analysis are on bottom. Dots represent median expression for individual samples. Line represents the mean for the four samples in each group. **(C)** Volcano plot of expression differences between TA and quadricep from FSHD biopsies from [26]. Genes with higher expression in TA or quadricep are in purple and green, respectively. FSHD-induced genes are colored in black. The number of genes higher in each muscle are labelled at the top. **(D)** Heatmap of FSHD-induced gene expression detectable in control quadricep, FSHD quadricep and FSHD TA biopsy samples from [26]. Groups were identified in [26] with an increase in group number correlating with an increase in DUX4 target gene expression. Clinical severity score (CSS) is on a 10-point scale. **(E)** Scatterplot of percent methylation in the promoters of FSHD-induced genes for control and FSHD2. Lines indicate a 25% difference. Promoters that overlap known DUX4 binding sites are colored orange. **(F)** Motifs from [35] or [64] enriched in promoters of FSHD-induced genes.

**Figure S4:**
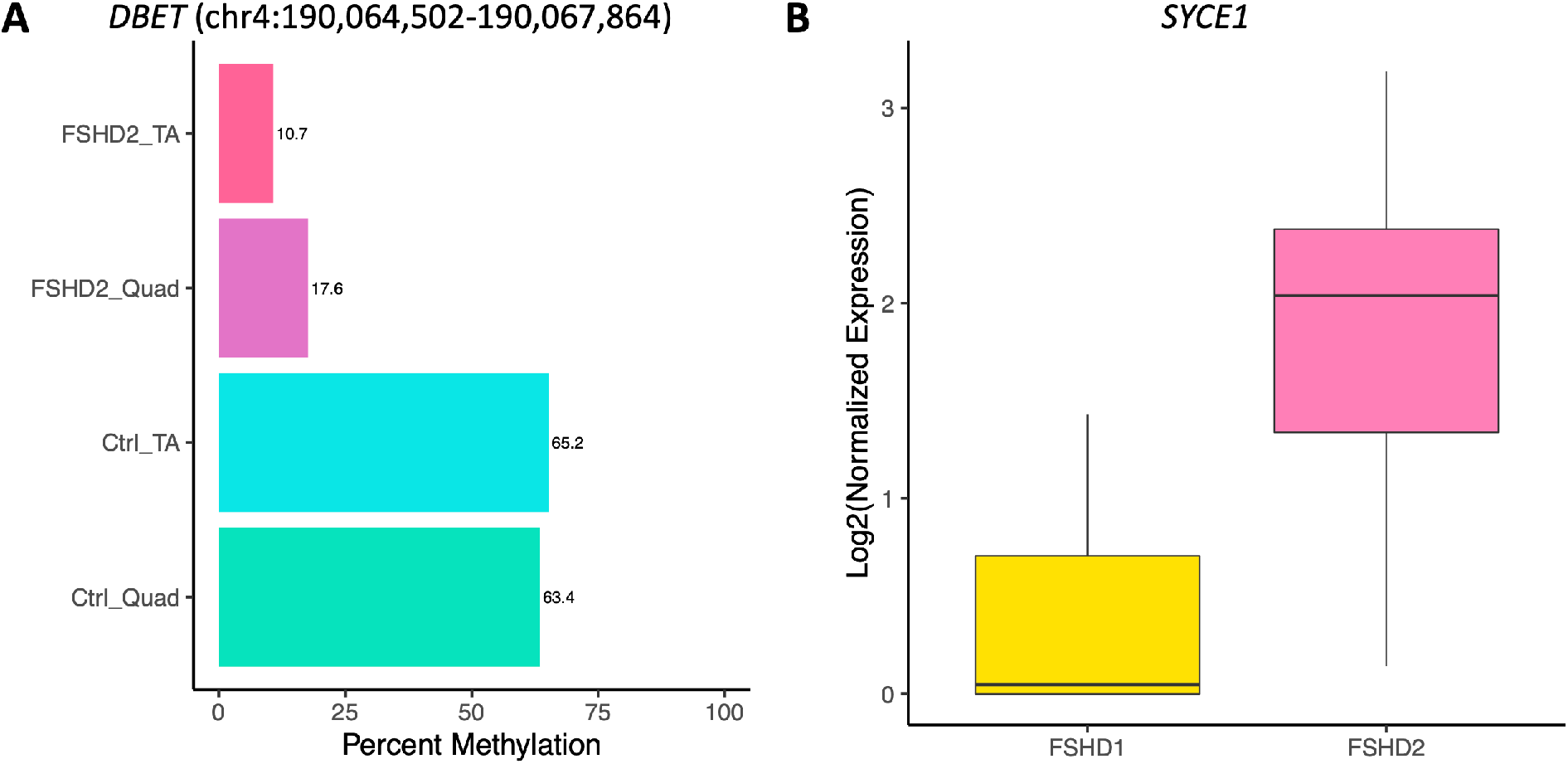
SMCHD1 related methylation and expression differences. **(A)** Barplot of average percent methylation near *DBET* on chromosome 4. Numbers to the left of bars give the average percent methylation. **(B)** Boxplot of *SYCE1* expression in FSHD1 and FSHD2 samples at day 0 of differentiation.

**Figure S5:**
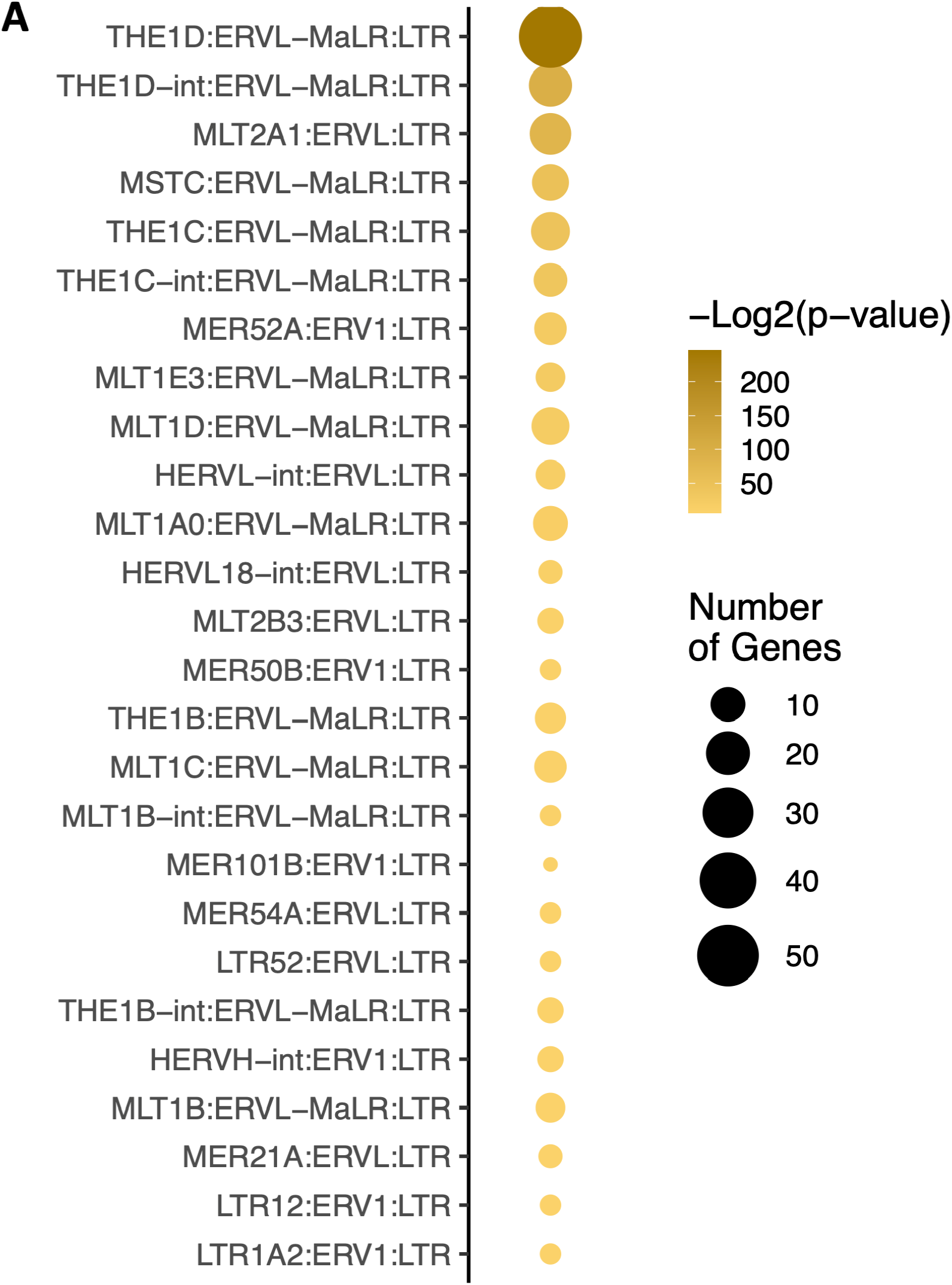
Enrichment of specific types of LTRs. **(A)** Enrichment of specific types of LTRs upregulated in FSHD2. Classes with p-value >0.05 are not shown. Size indicates number of loci in category.

**Figure S6:**
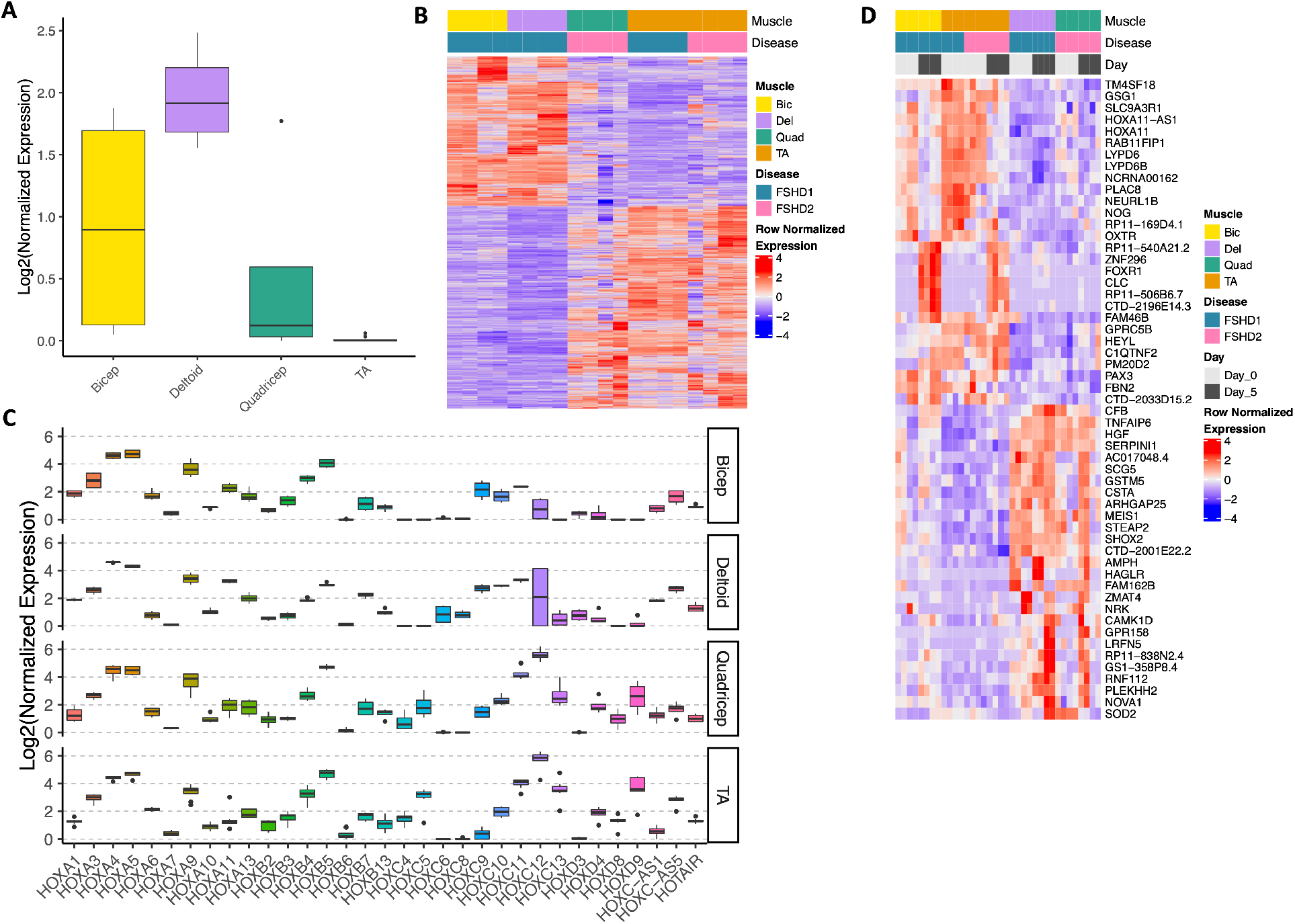
Muscle group specific expression. **(A)** Boxplot of *TBX5* expression in the different muscle groups at day 0 of differentiation. **(B)** Heatmap of genes specific to the upper (bicep and deltoid) or lower (TA and quadricep) body muscles at day 0 of differentiation. **(C)** Boxplot of HOX gene expression at day 0 of differentiation split by muscle group. **(D)** Heatmap of genes with differential expression in highly (TA and bicep) and less (quadricep and deltoid) susceptible muscle groups at days 0 and 5 of differentiation combined.

**Table S1: Differentially methylated regions between TA and quadricep in control cells** Genes are annotated by GREAT. Percent difference in methylation given in “meth.diff” column with positive values indicating higher methylation in TA.

**Table S2: Differentially expressed genes between TA and quadricep in control cells** TFs are labelled as well as TFs whose motifs were enriched in differentially methylated regions.

**Table S3: Motifs enriched in promoters of FSHD-induced genes**

**Table S4: Differentially methylated regions between FSHD2 and control** Genes are annotated by GREAT. Percent difference in methylation given in “meth.diff” column with positive values indicating higher methylation in FSHD2.

**Table S5: Differentially expressed genes between FSHD2 and control**

**Table S6: 580 TE loci with increased expression in FSHD2 during myogenesis** Loci that overlap DUX4 binding sites of FSHD-induced genes are indicated in the “DUX4_BS” and “OverlapFI” columns, respectively. Loci that overlap peaks from [24] are also indicated.

**Table S7: Genes with expression differences for highly susceptible and less susceptible muscle groups**

